# Engineering lung-sensing T cells using synthetic receptors targeting RAGE

**DOI:** 10.1101/2025.06.06.658315

**Authors:** Anjali Jacob, Caroline Whitty, Gian Luca Lupica-Tondo, Eshan S Badwal, Xin Ren, Wenli Qiu, Wei Yu, Yurie Tonai, Juan Antonio Camara Serrano, Milos S. Simic, Wendell A. Lim, Greg M. Allen, Dean Sheppard

**Affiliations:** Division of Pulmonary, Critical Care, Allergy and Sleep, Department of Medicine, University of California, San Francisco, San Francisco, CA, USA; UCSF Cell Design Institute and Department of Cellular & Molecular Pharmacology, University of California San Francisco, San Francisco, CA, USA; Department of Medicine, University of California San Francisco; San Francisco, CA, USA; USA Helen Diller Family Comprehensive Cancer Center, University of California San Francisco, San Francisco, CA, USA; Parker Institute for Cancer Immunotherapy, San Francisco, CA, USA

**Author notes:** Corresponding Author: Dean Sheppard PO BOX 589001, Room 252R University of California San Francisco San Francisco, CA 94158-9001.

## Abstract

Our goal was to leverage synthetic biology approaches to engineer lung-sensing T cells that trigger synthetic transcriptional programs only when in the lung. First, we identified a lung-specific cell surface protein, receptor for advanced glycation endproducts (RAGE), expressed at high levels exclusively in the lung. Then, we engineered chimeric antigen receptors (CAR) and synthetic notch receptors (SynNotch) to bind this protein. We showed that anti-RAGE CAR-T cells traffic to and proliferate in the lung exclusively. Anti-RAGE SynNotch receptors activate transcription of a fluorescent reporter only when co-cultured with RAGE+ cell lines or primary lung cells. Finally, we tested an anti-RAGE SynNotch to anti-CD19 CAR circuit in vivo in mice implanted with lung and flank tumors and found that this approach cleared CD19-expressing lung tumors without affecting CD19-expressing flank tumors. Thus, we demonstrate that RAGE is a lung-specific target and that T cells expressing anti-RAGE receptors can sense and activate lung-specific therapeutic transcriptional programs. This approach could be extended to allow cell-based therapies that target and deliver genetically encoded payloads to treat lung diseases while avoiding systemic toxicity.

## Introduction

Systemic administration of drugs and cell-based therapies is fraught with off-tissue, on-target toxicities. For diseases primarily confined to one tissue, strategies that allow for delivering therapies only to a specific tissue would avoid toxicities in unrelated tissues. Prior work has shown that T cell therapies can be programmed to recognize brain-specific antigens and locally execute therapeutic functions only within the brain, including tumor killing^1^ or suppression of inflammation.^2^ To extend the utility of tissue-specific cell-based therapies to non-CNS tumors and diseases, we need to develop recognition programs for other organs.

Lung diseases have historically had few available mechanism-specific therapies. Lung tumors are primarily treated surgically, sometimes requiring the removal of large areas of healthy lung, which can be challenging in patients who are elderly or have many co-morbidities. Highly morbid conditions affecting the lung, like acute respiratory distress syndrome (ARDS) and pulmonary fibrosis, have no therapies with a mortality benefit^3^. Though cell-based therapies, specifically mesenchymal stromal cells, have been explored for ARDS, the mechanism by which they confer protection in preclinical models is unclear, and they have not thus far shown benefit in clinical trials^4^. T cells have been used extensively in engineered cell-based therapies, primarily for oncologic purposes, but recently they have been considered for treating autoimmune and fibrotic diseases.^5,6^ We aim to extend cell-based therapy technologies to treat diseases that affect the lungs.

We developed a plan to engineer T cells to recognize a cell surface protein highly expressed in the healthy lung but minimally expressed in other tissues. We used bioinformatic analysis to identify the receptor for advanced glycation endproducts (RAGE) as the most promising target to pursue^7,8^. We developed a chimeric antigen receptor that targets RAGE and found that after adoptive transfer into mice, RAGE-CAR T cells traffic to and proliferate in the lung where they cause local lung inflammation, confirming that this is an effective lung targeting technique. We also developed a synthetic notch (SynNotch) receptor that recognizes RAGE and showed that when mice that harbor tumors in their lungs and flanks are treated with T cells containing a RAGE-SynNotch to a tumor-killing CAR circuit, only the lung tumors are controlled, and the flank tumors continue to grow. Thus, we have developed novel receptors targeting a lung-specific surface protein that enable the programming of lung-specific T cell behaviors that will have practical applications in treating lung disease.

## Results

### RAGE is a lung-specific surface protein that could be used as a target for recognition by CAR and SynNotch receptors

Cell surface receptors enable T cells to recognize and respond to their environment. Our goal was to engineer synthetic receptors that allow T cells to activate transcriptional programs when they are in the lung (Fig. 1A). First, we sought a surface protein that was expressed specifically in the lung. The lung’s most differentially expressed surface gene compared to other tissues was AGER, or advanced glycation end-product receptor, which encodes the protein RAGE, receptor for advanced glycation end-products.^9^ RAGE is expressed in type I alveolar epithelial cells (AT1s), which are the large, flat cells over which oxygen and carbon dioxide are passively exchanged with pulmonary capillary endothelial cells in the alveoli^8,10^ (Fig. 1B, C). Because AT1s cover 95% of the lung’s surface area, they were an attractive target for lung-sensing T cells, since T cells are more likely to encounter RAGE+ AT1s than any other lung epithelial or interstitial cell type. We analyzed GTEx data and found that AGER is expressed highly in the lung and at low levels in the thyroid but is minimally expressed in other tissues^9^ (Fig. 1D). Single-cell RNA sequencing analysis confirms that AGER is highly expressed in AT1s, with lower-level expression in AT2s (Fig. 1E). Importantly, prior work has shown that RAGE is expressed in the context of lung disease, both in acute lung injury and in the transitional AT2 cells that arise in the context of pulmonary fibrosis^8,11^ (Fig. 1E).

**Figure 1.**
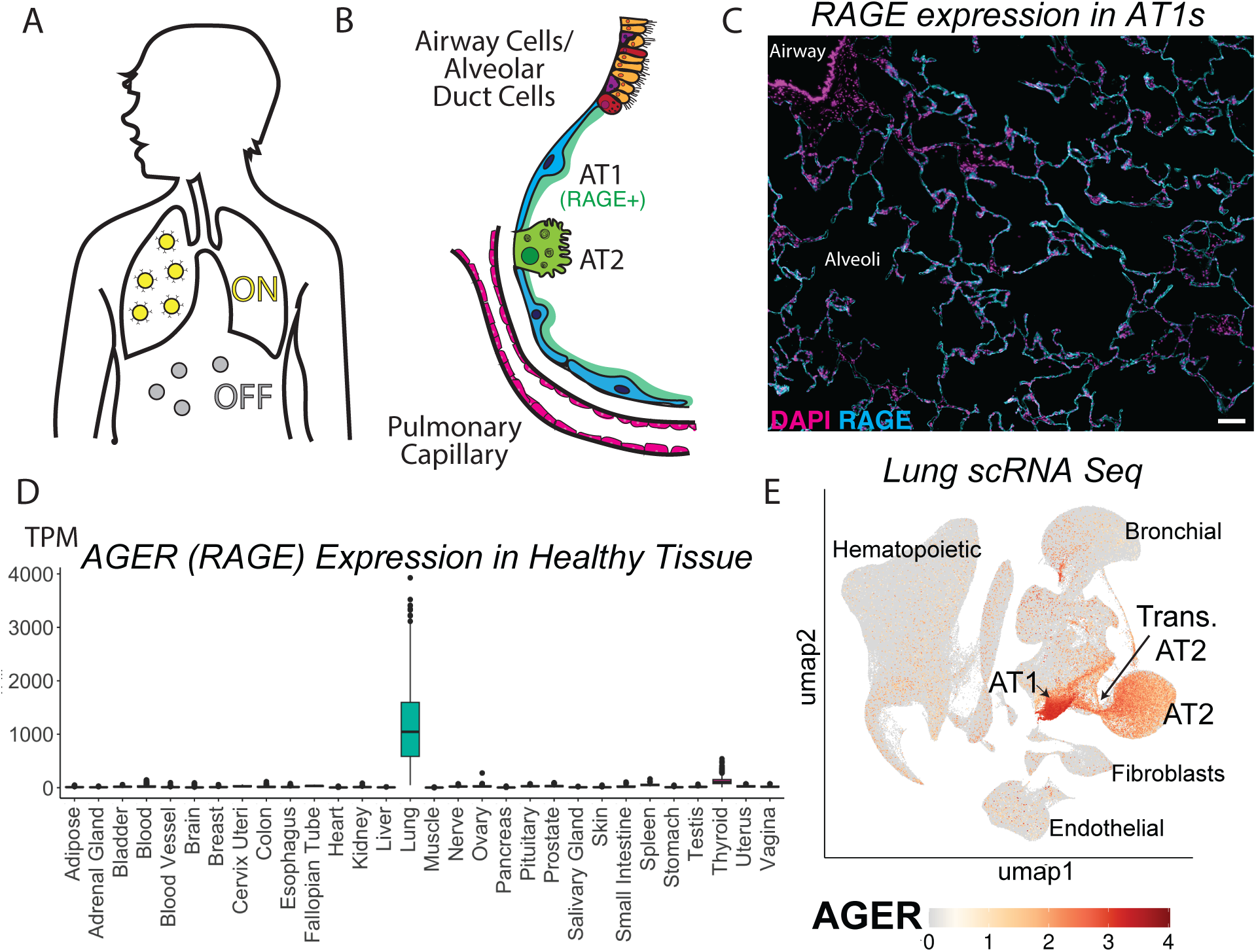
Rationale for RAGE as a lung specific target. A) Schematic showing conceptual goal for T cells to turn on a programmed circuit when they sense the lung, while it remains off in other tissues. B) Cellular architecture of the distal lung with RAGE expressing alveolar type I cells (AT1s) in blue that cover the majority (95%) of the alveolar surface area. Alveolar type II cells (AT2), other epithelial cells (airway/alveolar duct), and endothelial cells in pulmonary capillaries also noted. C) Immunofluorescence of cell nuclei (DAPI, magenta) and RAGE (cyan) in healthy human lung tissue with airways noted to be RAGE negative and the majority of the alveolar space surface area noted to be RAGE positive. Scale bar 100um. D) GTEx data showing AGER (gene encoding RAGE) expression by tissue. E) Single cell RNA-sequencing data from Natri et al, Nature Genetics 2024 showing AGER expression in single cells from both pulmonary fibrosis and healthy human lungs. AGER is highest in AT1s but also expressed at lower levels in AT2s and transitional cells (damaged alveolar epithelial cells that appear in the context of pulmonary fibrosis)

### Anti-RAGE CAR T cells traffic to the lung and cause lung injury

We used the ScFv from a published soluble-RAGE targeting antibody that binds to the extracellular domain of cell surface RAGE to generate a chimeric antigen receptor.^12^ In vitro, these anti-RAGE CAR T cells kill K562s engineered to overexpress RAGE (Fig. 2A, B).

**Figure 2.**
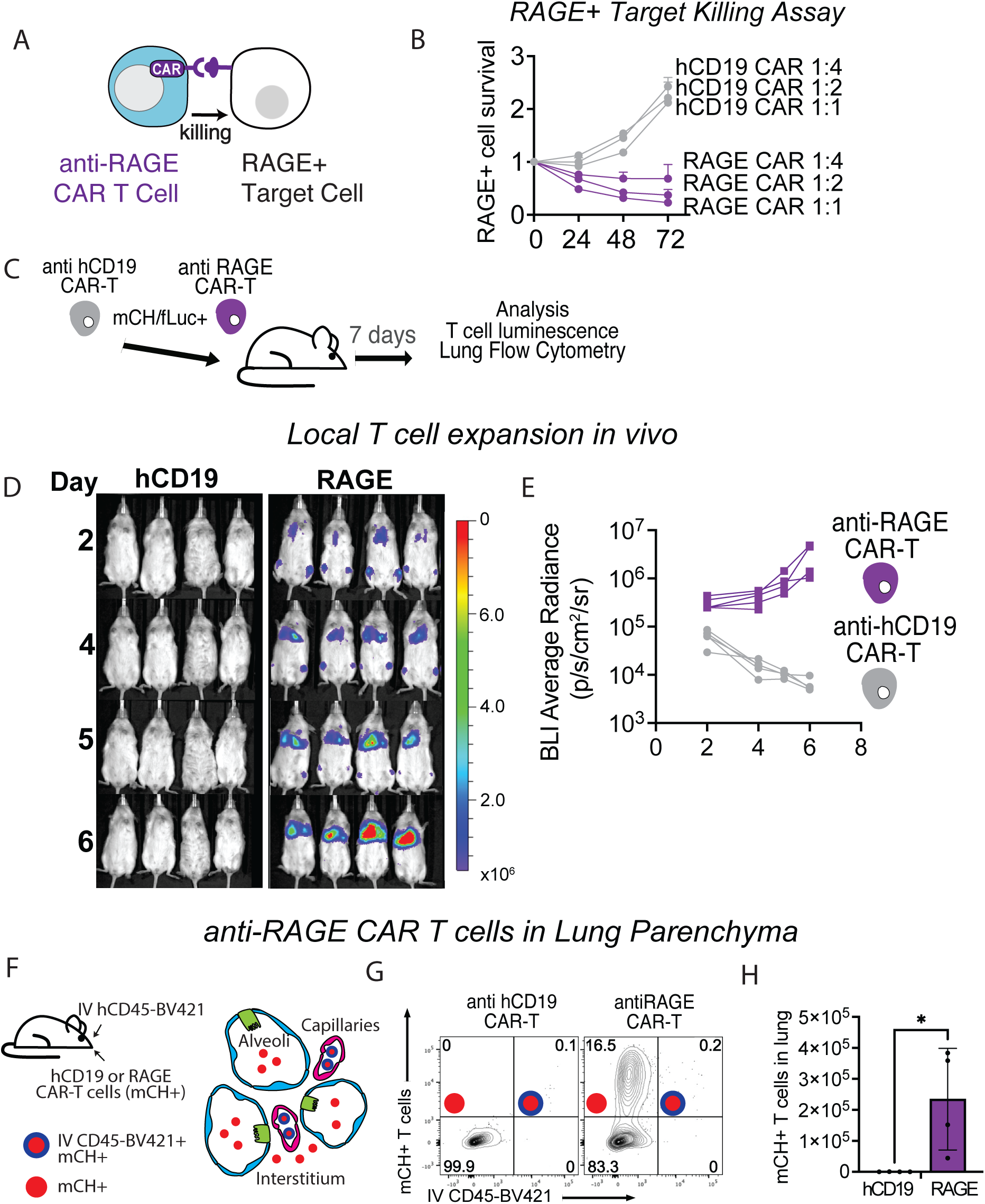
anti-RAGE CAR enables T cells trafficking to and proliferation in the lung. A) Schematic showing concept of in vitro killing assay wherin antiRAGE CAR T cells kill K562 engineered to overexpress RAGE. B) Fold change in live RAGE expressing K562s when cultured with anti-human CD19 CAR-T cells or anti RAGE CAR-T cells. Ratios are shown as CAR-T cell (effector): K562 (target). C) Schematic showing in vivo experiment where anti-human CD19 CAR-T cells were compared to anti RAGE CAR-T cells when injected via tail vein into NSG mice. CAR-T cells were cotransduced to express mCherry and luciferase constitutively. D) 6 day time course of in vivo bioluminescence of luciferase+ T cells. E) Quantification of luminescence in lung fields over time from figure 2D. F) Schematic showing concept behind IV human CD45 experiment. Mice injected with IV CD45 have intravascular T cells labeled with IV human CD45 and extravascular T cells unlabeled. Thus, T cells in the lung parenchyma are mCherry+ alone while T cells in the pulmonary vasculature are IV human CD45-BV421+ and mCherry+. G) IV human CD45 BV421 and mCherry flow cytometry data from mouse lungs digested on Day 7 after injection with CAR-T cells showing very few mCH+ cells in the anti human CD19 CAR-T cell group and primarily mCH+ BV421-cells in the anti RAGE CAR-T cell group suggesting T cells are primarily extravascular in the lung parenchyma. H) Quantification of total mCH+ T cell number in left lung of each mouse.

To evaluate how specific anti-RAGE CAR T cells would be for lung targeting in vivo, we generated mCherry+ f-luciferase+ anti-RAGE CAR T cells and delivered them via tail vein injection into NSG mice. We compared these mice to mice receiving anti-human CD19 CAR T cells that could have some tonic CAR activity but do not target any endogenously expressed protein in the mouse (Fig. 2C). We found that within days, luminescence in the lung fields of mice receiving anti-RAGE CAR T cells was present at higher levels than in mice receiving control anti-hCD19 CAR T cells and that lung field luminescence in mice receiving anti-RAGE CAR T cells increased over time while lung field luminescence in the mice receiving anti-hCD19 CAR T cells decreased over time (Fig. 2D, E). Ex vivo imaging of other organs showed that the T cell luminescence in mice receiving anti-RAGE CAR-T cells was seen primarily in the lungs and not in any other thoracic organs to explain the luminescent signal in the in vivo images (Fig. S2A).

Immune cells are frequently found in the lung’s intravascular spaces, even when activated in other body regions, because the entire blood volume circulates through the lung for gas exchange. To determine whether anti-RAGE CAR T cells were present only intravascularly or actually in the lung parenchyma, we injected mice with IV human CD45-BV421 antibody and euthanized and harvested their lungs 3 minutes later, a technique that has been shown to label all intravascular cells with antibody, but no parenchymal cells^13^. Since CAR T cells were co-transduced with mCherry, we could determine whether T cells were intravascular (BV421+, mCH+) or parenchymal (BV421-mCH+) (Fig. 2F). We found that there were minimal mCH+ T cells in the lungs of mice that receive anti-hCD19 CAR T cells and almost all of the mCH+ T cells in the mice that received anti-RAGE CAR T cell were BV421-, indicating that they were indeed in the lung parenchyma (Fig. 2G, H).

Finally, we assessed whether anti-RAGE CAR T cells were causing lung inflammation or damage. In healthy lungs, BAL fluid has no neutrophils, but when lung inflammation leads to alveolar damage, the typically tight alveolar membranes become leaky and neutrophils extravasate into the alveolar space^14^. Bronchoalveolar lavage (BAL) fluid from mice receiving anti-RAGE CAR T cells showed an increased number of neutrophils in the lavage fluid, suggesting that there was alveolar damage in the mice that received anti-RAGE CAR T cells (Fig. S2B).

### Engineering an anti-RAGE SynNotch receptor for lung sensing

Having validated the strategy of using RAGE as a lung-specific target, we next aimed to engineer a receptor that would enable T cells to express a genetically encoded payload (eventually a therapeutic) upon sensing lung-specific RAGE without the cell-mediated killing of important alveolar epithelial cells and lung inflammation caused by a CAR. We used the same anti-RAGE ScFv^12^ as the recognition domain of a synthetic notch receptor. We showed that when T cells expressing an anti-RAGE SynNotch to BFP circuit are cultured with mouse or human RAGE+ K562s, mouse lung cells in vitro, or precision-cut lung slices that preserve the architecture of the mouse lung while being cultured in vitro^15^, T cells do activate expression of their fluorescent reporter (Fig. 3A-D). Thus, the anti-RAGE SynNotch enables T cells to sense the lung via RAGE.

**Figure 3.**
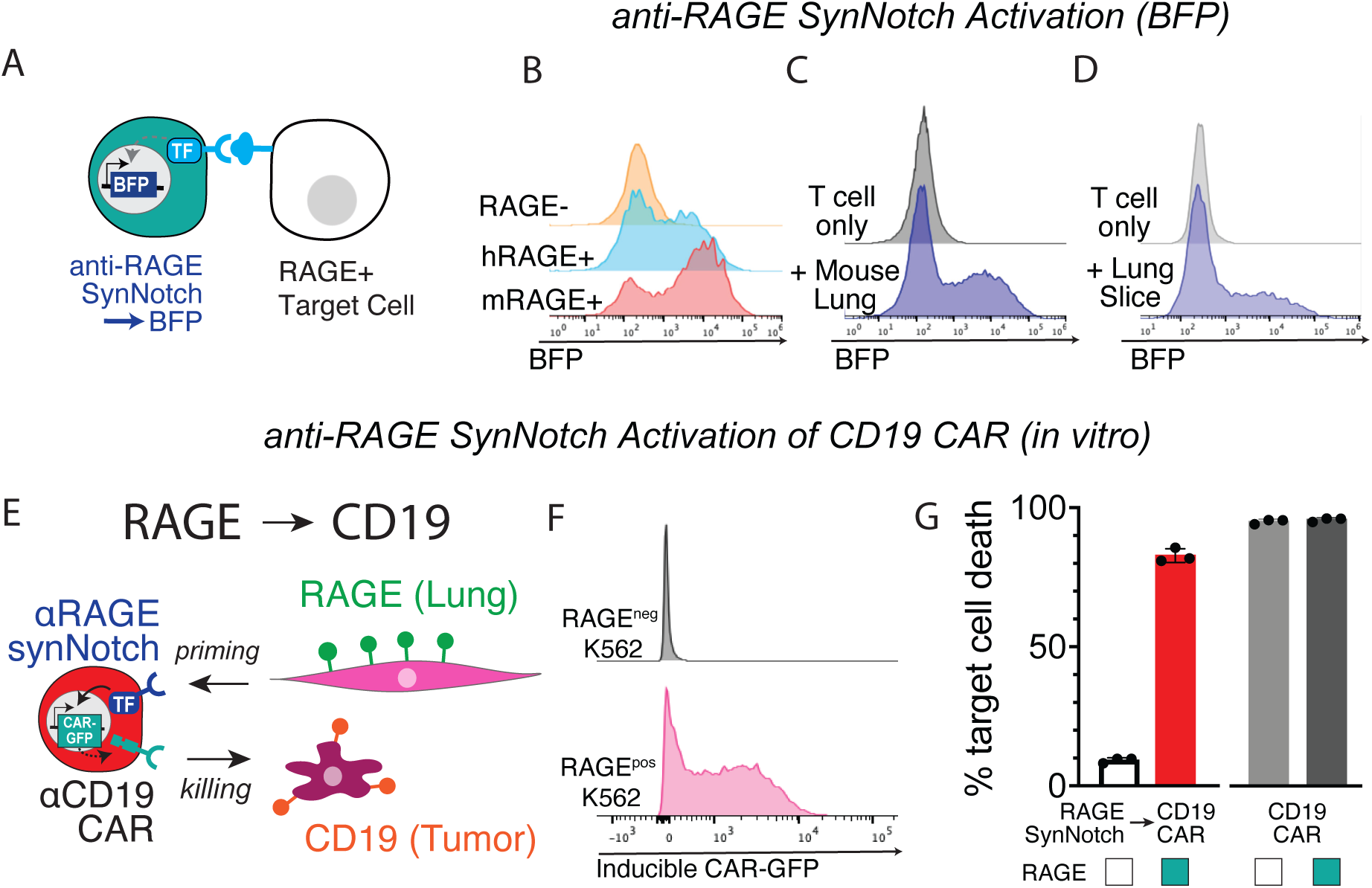
T cells can turn on gene expression in response to anti-RAGE SynNotch receptor recognition of RAGE+ cells and lung cells in vitro. A) Schematic showing anti RAGE SynNotch-to-BFP circuit in T cells in which T cells should express BFP when cultured with RAGE+ cells. T cells in Fig. 3B-D express this circuit. B) BFP expression in T cells when cultured with RAGE-K562 or K562 that express the human form of RAGE or the mouse form of RAGE. C) BFP expression in T cells when cultured with mouse lung cells. D) BFP expression in T cells when cultured with precision-cut lung slices from healthy mouse lungs. E) Schematic showing anti RAGE SynNotch-to-CD19 CAR-2a-GFP circuit in which T cells should express CD19 CAR and GFP when cultured with RAGE+ cells. F) GFP expression in T cells with the RAGE SynNotch to CD19 CAR-2a-GFP circuit when cultured with RAGE-K562 (top) or RAGE+ K562 (bottom). G) Target cell death after 72 hours when CD19+ NIH1975 tumor cells are cultured with RAGE SynNotch to CD19 CAR circuit T cells and RAGE-K562s or RAGE+ K562s, compared to constitutively active CD19 CAR-T cells.

### Anti-RAGE synNotch → CD19 CAR circuit causes CD19+ tumor killing in vitro only in the presence of RAGE+ cells

As a proof-of-concept to test if the anti-RAGE synNotch receptor could be used to activate cellular function in the lung with a measurable therapeutic response, we designed T cells expressing the anti-RAGE synNotch receptor, along with a synNotch responsive promoter that induces the expression of an anti-human CD19 CAR-2a-GFP. In principle, this circuit would yield lung-sensing T cells that would activate a tumor-killing CAR only in the lung (Fig. 3E). Indeed, in vitro T cells expressing the anti-RAGE SynNotch to CD19 CAR-GFP only expressed CAR-GFP when cultured with RAGE+ K562s, not RAGE-K562s. These T cells could kill tumor cells expressing CD19 to a similar degree as T cells that constitutively expressed an anti-CD19 CAR after 72 hours, but only when cultured with RAGE+ K562s (Fig. 3F, G).

### Anti-RAGE SynNotch →CD19 CAR T cell clears tumors implanted in the lung but not outside the lung

We next sought to test the anti-RAGE SynNotch in a mouse model where tumors were implanted simultaneously into the lungs and flanks of mice. We selected a tumor line that grew similarly in the lung and flank regions of NSG mice, NCI-H1975 lung adenocarcinoma, engineered to overexpress human CD19+. Because we wanted to test whether the anti-RAGE SynNotch was sufficient to induce tumor killing in the lung, we chose to use a well-validated CAR T cell target, human CD19, rather than an endogenous tumor target. In the absence of treatment, the lung tumor alone led to mortality within 24-30 days (Fig. 4D). Mice harboring both a lung and flank tumor were divided into 3 groups and treated with T cells either 1) untransduced and not expressing a CAR (UnT); 2) expressing an anti-human CD19 CAR under the control of an anti-RAGE SynNotch receptor, thus CAR predicted only to be expressed in the lung (Lung primed CAR) or 3) constitutively active human CD19 CAR T cells active in all locations in the body (CAR, Fig. 4A-C). Because lung tumors could not be measured directly, we heterologously expressed luciferase in the tumor cells and measured luminescence of lung and flank tumors. In the UnT group, neither lung nor flank tumors were controlled. In the CAR group, both lung and flank tumors were controlled. In the SN group, the lung tumors were controlled, but flank tumors persisted, confirming that the anti-RAGE SynNotch enables lung-specific gene expression, in this case of the anti-hCD19 CAR (Fig. 4E, F; S3A-D). These data confirmed that we could use the anti-RAGE SynNotch to turn on genetic circuits specifically in T cells in the lung without off-target effects at other sites.

**Figure 4.**
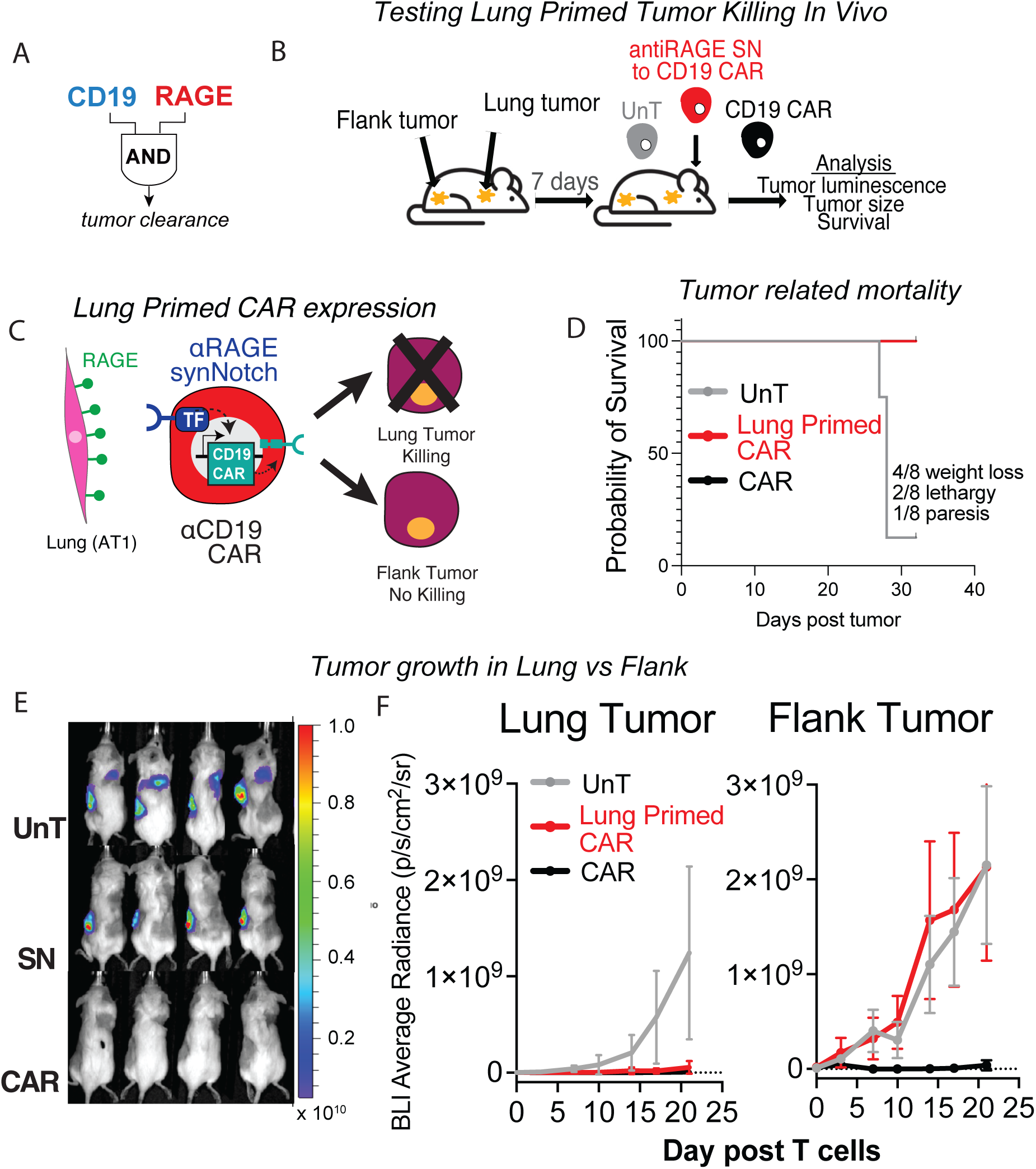
anti-RAGE SynNotch →CD19 CAR enables CD19 expressing tumor clearance specifically in the lung. A) Concept that only when T cells recognize both RAGE and CD19 will target cell killing occur. B) Schematic of experiment in which mice were injected with tumors (NIH1975 lung cancer overexpressing hCD19) in both lung and flank and treated with T cells 7 days later. Mice were split into 3 groups: untransduced T cells (UnT), anti-RAGE SynNotch to hCD19 CAR circuit (SN), and constitutive CD19 CAR (CAR) and mouse survival was tracked over time, tumor burden was analyzed via in-vivo bioluminescence. C) Schematic of expected results from this experiment. In the lung, T cells would recognize RAGE, activating transcription of the CD19 CAR, leading to tumor killing whereas in the flank, T cells would not recognize RAGE, not activate transcription of the CD19 CAR, and tumors would persist. D) Survival curves show 7/8 mice in UnT group die within 31 days, while all mice in the SN and CAR groups are alive at this timepoint. P<.0001 by Mantel-Cox regression. E) In vivo bioluminescence of luciferase+ tumor in UnT, SN, and CAR groups on day 31 post tumor injection (day 24 post T cell injection). Lung and flank luminescence is visible in UnT group, flank only in SN group, neither in CAR group. F) Quantification of luminescence in lung fields and flanks in mice from each group. Error bars represent standard deviation.

## Discussion

### A platform for engineering therapeutic cells that target their activity selectively to the lung

We report the generation of anti-RAGE receptors that enable T cells to sense that they are in the lung and allow programming of further behaviors, like CAR-mediated tumor killing, to be restricted to the lung. This technology will be useful in programming T cells to deliver a variety of therapeutics to the lungs. Our group has previously used brain-specific T cell circuits to deliver CAR T cells that target primary or metastatic brain tumors and control tumors in the brain without being active in other tissues^1,2^. We have also used brain-specific SynNotch expressing T cells to deliver anti-inflammatory cytokines to the CNS, ameliorating autoimmunity in mouse models of multiple sclerosis^2^. Similarly, by varying the payloads controlled by anti-RAGE SynNotch binding, we could restrict expression of molecules that could be useful in treating lung diseases to the lung, concentrating their expression in the target tissue, and limiting off-target effects. Anti-lung tumor payloads could more potently control lung tumors, anti-inflammatory payloads could treat autoimmune lung diseases, anti-fibrotic payloads could treat pulmonary fibrosis and interstitial lung diseases, and growth factor payloads could improve lung regeneration after injury. Using engineered cells to deliver signaling pathway modulators to the lung would also enable a temporally regulated study of how signaling pathway activation or inhibition affects the progression of lung disease without using laborious genetic mouse models.

The surface area of the human lung makes it virtually impossible to intratracheally deliver cell-based therapies that would reach the most distal regions of the lung alveolus, which are often affected by incurable lung diseases like idiopathic pulmonary fibrosis or acute respiratory distress syndrome (ARDS).^16^ Intravenously injected drugs traffic first through the pulmonary vasculature but transit into other tissues and can cause on-target, off-tissue effects. Using anti-RAGE CAR and SynNotch receptors would allow for cell-based therapies administered intravenously to have lung-specific therapeutic effects. Anti-RAGE CAR T effector cells are likely too toxic to be clinically useful. Still, the anti-RAGE CAR could be used in regulatory T cells, which do not generally induce tissue injury and, even in the absence of CARs, have been shown to improve recovery in models of acute lung injury^17,18^. Anti-RAGE SynNotch T cells would enable cell-based therapies to be delivered systemically, but only activated locally in the lung.

New tools for understanding the biology and pathobiology of the lung and new mechanism-specific therapies for lung diseases are needed, and this work represents a novel synthetic biology approach to further our understanding of lung disease.

## MATERIALS AND METHODS

### Receptor Design

CAR receptors were built by fusing scFv sequences from patent to the hinge region of the human CD8a chain and transmembrane and cytoplasmic regions of the human 4-1BB, and CD3z signaling domains. SynNotch receptors were built by fusing scFv sequences to mouse Notch1 (NM_008714) minimal regulatory region (residues 1427 to 1752) and Gal4 DBDVP64. All anti-RAGE synNotch and CAR receptors contain N-terminal CD8a signal peptide (MALPVTALLLPLALLLHAARP) for membrane targeting and a-myc-tag (EQKLISEEDL) for detecting surface expression with a-myc A647 (Cell Signaling Technology, catalog no. 2233). See Morsut et al.^19^ for the synNotch sequence. Receptors were cloned into a modified pHR vector containing a PGK promoter. The pHR vector was also used to make response element plasmids with five copies of the Gal4 DNA-binding domain target sequence (GGAGCACTGTCCTCCGAACG) upstream from a minimal CMV promoter. Response element plasmids also contain a PGK promoter that constitutively drives mCH expression to identify transduced T cells easily. Inducible CAR constructs (anti-human CD19-GFP) were cloned into a site 3’ to the Gal4 response elements and minimal CMV promoter.

### T cell isolation and culture

T cells from human donors were isolated using a magnetic bead-based CD4 or CD8 isolation kit (StemCell Technologies). Blood was obtained from StemExpress or Allcells, as approved by the University of California, San Francisco (UCSF) institutional review board. T cells were cryopreserved in Cellbanker 1 or RPMI-1640 with 20% human AB serum (Valley Biomedical) and 10% dimethyl sulfoxide. After thawing, T cells were cultured in human T cell medium consisting of X-VIVO 15 (Lonza), 5% human AB serum, 55 µMb-mercaptoethanol, and 10 mM neutralized N-acetyl-L-cysteine supplemented with 30 units/ml IL-2.

### Lentiviral Transduction of T cells

For in vitro CAR and SynNotch experiments, CD8+ T cells (CAR) or CD4+ T cells (SynNotch) were transduced with lentiviral vectors. For in vivo CAR and SynNotch → CAR circuit experiments, CD8+ T cells and CD4+ T cells were transduced, and equal numbers were cultured separately until the day of T cell injection into mice, when in each experiment, they were injected at a 1:1 ratio. Pantropic vesicular stomatitis virus G (VSV-G) pseudotyped lentivirus was produced through transfection of Lenti-X 293T cells (Takara Bio, catalog no. 632180) with a pHR transgene expression vector and the viral packaging plasmids pCMV and pMD2.G using Fugene HD (Promega) or TransIT-VirusGen (Mirus Bio). Primary T cells were thawed the same day and, after 24 hours in culture, were stimulated with 25 µl of anti-CD3/CD28 coated beads [Dynabeads Human T-Activator CD3/ CD28 (Gibco)] per 1 ×10^6^ T cells. At 48 hours, viral supernatant was harvested. Primary T cells were spinoculated at 2000g for 2 hours on retronectin-coated plates. On days 5-6 after T cell stimulation, Dynabeads were removed, and T cells were sorted using a BD Biosciences FACSARIA Fusion. T cells were expanded until rested for 6-8 days before being used in assays.

### Cell lines

Cell lines used were K562 myelogenous leukemia cells (ATCC catalog no. CCL-243) and NCI-H1975 (ATCC catalog number CRL-5908). K562s were lentivirally transduced to express human or mouse isoforms of RAGE-2a-mCherry. NCI-H1975 cells were transduced using two vectors to express human CD19 and mCherry-luciferase. Cell lines were cultured in DMEM + 10% FBS.

### In vitro assays for anti-RAGE CAR function

CD8+ T cells were transduced with plasmids encoding anti-RAGE CAR or anti-human CD19 CAR, sorted 5 days after transduction, rested for 7 days, and cocultured with RAGE+ K562s 7 days after sorting. 24-72 hours after coculture, cells were harvested for flow cytometry and stained with CD8-GFP and Draq7. CD8-cells in the RAGE CAR experiment were K562s, and the total number of live K562 (CD8-/Draq7-) was counted each day and compared to the initial number seeded (fold change vs Day 0).

### Lung flow cytometry

For flow cytometry, mice were injected with retro-orbital anti-human CD45-BV421 (Biolegend) 3 minutes before euthanizing with isoflurane. Left lungs were minced and digested with collagenase I, dispase, and DNAse. Total cell suspensions were resuspended in 2% FBS in PBS (FACS buffer) and analyzed for BV421 and mCherry expression. Cell counts were obtained using CountBright flow cytometry counting beads.

### Mouse lung cell isolation and culture

Mouse lung cells were dissociated as described above and EPCAM+ MHCII+ sorted cells from C57/Bl6 mice grown in 2D culture. 20K cells were plated into each well of a 96-well plate culture in DMEM + 10% FBS for 5 days.

### Mouse precision cut lung slice

Lungs from healthy C57/Bl6 mice were inflated with 2% low melting temperature agarose and removed after mice were euthanized. After 20-30 minutes on ice, we glued one lobe of the lung at a time to the stage of a Leica VT1200 vibratome and cut 300 µm thick sections at 0.70 mm/s. Sections were placed into a 24 well plate in PBS on ice.

Lung slices were then transferred into new 24 well plate with anti-RAGE SynNotch T cells in human T cell media (described above). Lung slices were dissociated as described above and analyzed by flow cytometry, with T cells stained for CD3 and flow cytometry showing BFP expression in CD3+ population.

### In vitro assays for anti-RAGE SynNotch function

CD4+ T cells were transduced as above with plasmids encoding anti-RAGE SynNotch were sorted 5 days after transduction, rested for 7 days, and cocultured with mouse or human RAGE+ K562s, mouse lung cells, or precision cut mouse lung slices 7 days after sorting. 48 hours after co-culture, T cells were analyzed for BFP expression by flow cytometry.

To test the anti-RAGE SynNotch → anti-human CD19 CAR circuit, CD8+ T cells were transduced as above with plasmids encoding anti-RAGE SynNotch → anti-human CD19 CAR-GFP circuit. They were sorted 5 days after transduction, rested for 7 days, and cocultured with mouse RAGE+ or parental K562s and NCI-H1975 tumor cells transduced with human CD19 at a 1:1:1 (T cell:K562:Tumor cell) ratio. CAR-GFP expression in live T cells was analyzed, and NCI-H1975 cells were stained with Celltrace violet before the start of the experiment. Percent target cell death was determined for live NCI-H1975 cells as percent Draq7+ Celltrace violet+/ total Celltrace violet+ cells.

### Adoptive transfer of T cells

T cells from human donors were isolated using a magnetic bead-based CD4 or CD8 isolation kit (StemCell Technologies). For anti-RAGE CAR experiments, CD8+ T cells and CD4+ T cells were transduced with lentiviral vectors encoding anti-RAGE CAR, luciferase, and mCherry. 4 days after transduction, they were sorted, and 7 days after sorting, 2e6 cells (1e6 CD4 and 1e6 CD8) were injected into the tail vein of NSG mice.

For anti-RAGE SynNotch → CD19 CAR experiments, CD4 and CD8 T cells were prepared as described above in 3 groups: Untransduced T cells, T cells transduced with anti-RAGE SynNotch and anti-human CD19 CAR response element, and T cells transduced to express the same anti-hCD19 CAR constitutively. 7 days after tumor injections, mice were intravenously injected (retro-orbital) with 6e6 total (3e6 CD4 and 3e6 CD8) T cells from each group. The experiment was concluded after 31 days. At the endpoint, lungs were harvested and embedded in paraffin, and flank tumors were harvested.

### Bronchoalveolar Lavage

Bronchoalveolar lavage was performed by instilling 1 ml of PBS into the mouse lungs 3 times. Cells were harvested by centrifugation, resuspended in formalin, cytospun to adhere cells to slides, stained with hematoxylin and eosin, and lymphocytes, neutrophils, monocytes, and eosinophils were counted.

### Tumor injection in vivo

1e6 hCD19+ mCH+ fLuc+ NIH1975 cells were injected into the right lungs and left flanks of NSG mice. For lung injection, mice were anesthetized with isoflurane and placed on their left side, with the right chest region shaved and cleaned with iodine. 1e6 tumor cells were resuspended in 50 μL PBS and injected via a 30 G needle into the chest space of the mice at a depth of 7-8 mm. To ensure that intrathoracic vital structures were not damaged, the right front limb was moved back until the paw reached the ventral aspect of the thorax, with the olecranon marking the anatomical reference for the injection point. Injection was done during inspiration and paused during expiration, and the needle was kept in place for 30-60 s to avoid leakage of tumor cells into the extrapulmonary space. In 10% of mice, extrapulmonary tumors were noted in the subcutaneous area around the injection site, and these mice were excluded from further analysis. For flank tumors, the left flank region of the mice was shaved and cleaned with iodine, and 1e6 of the same tumor cells injected into the lung were injected subcutaneously immediately after lung tumor injection.

### Bioluminescence analysis

Live mice were injected with 3 μg luciferin, and live mice or dissected organs were analyzed on Xenogen IVIS 2000 10 minutes after 3.0 mg of D-luciferin (GoldBio, injection volume 200 μL) for CAR-T cell experiments and 1.5mg for tumor experiments. For tumor experiments, before treatment, mice were randomized so that the initial tumor burden in the control and treatment groups was equivalent. For flank tumor analysis, mice were imaged in a prone position. For lung tumor analysis, mice were imaged with right side up and a black barrier covering the lower half of the body to avoid bleed-through of more superficial flank signal. Average radiance was calculated for the lung and flank fields.

### In vivo experiments

Mice were monitored either daily (CAR) or twice weekly (SN→ CAR), and survival was evaluated over time until a predetermined IACUC-approved endpoint (e.g., tumor size, hunching, respiratory distress, paralysis, weight loss) was reached.

## Supplementary Figures

**Supplementary Figure 1.**
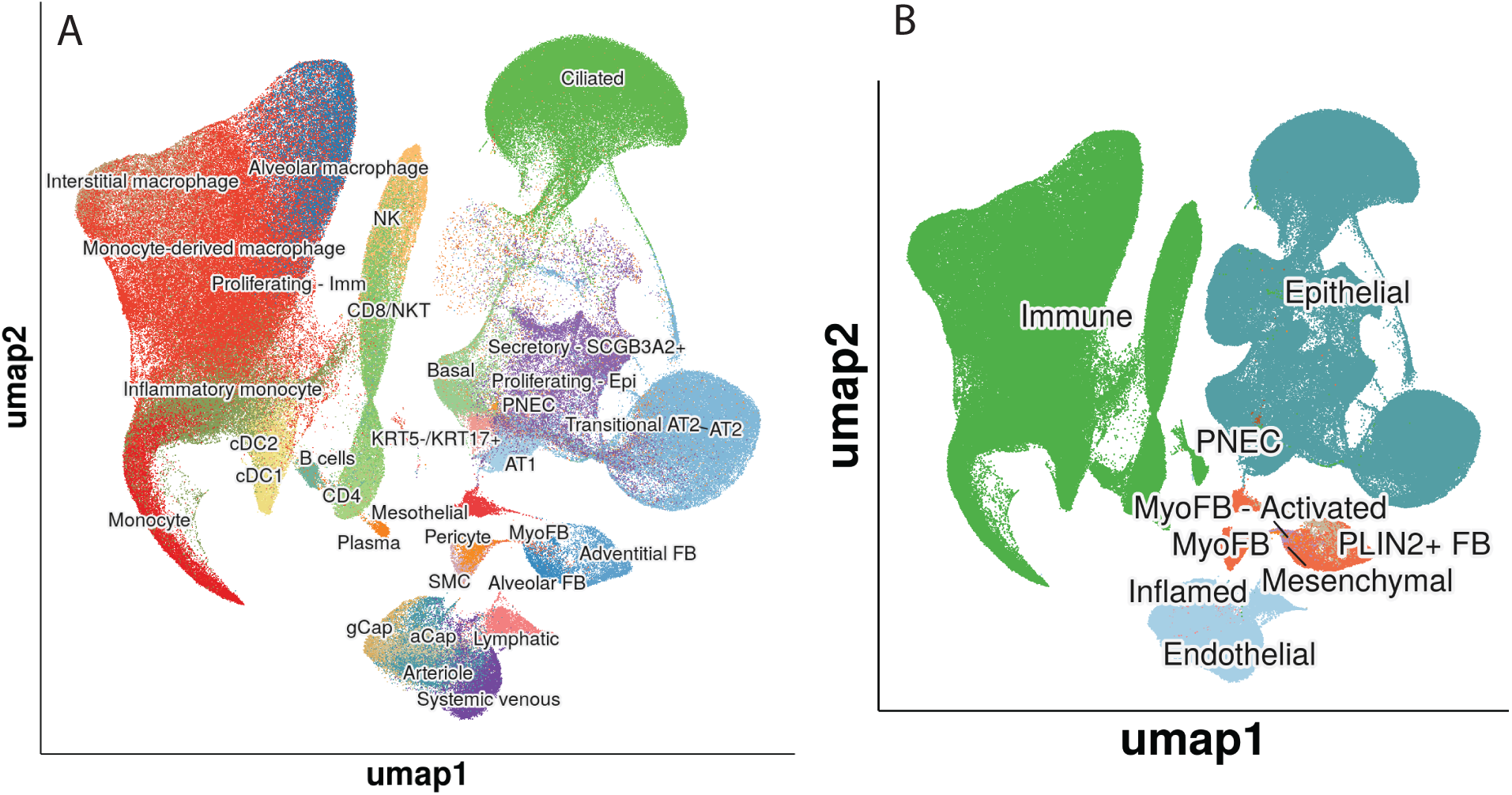
Single Cell RNA Sequencing annotations from healthy and IPF lung. Single cell RNA-sequencing data from Natri et al, Nature Genetics 2024 with A) manual annotations and B) lineage specific annotations directly obtained from https://app.lungmap.net/app/shinycell-ild-natri-2024. These annotations are partially represented in Figure 1E.

**Supplementary Figure 2.**
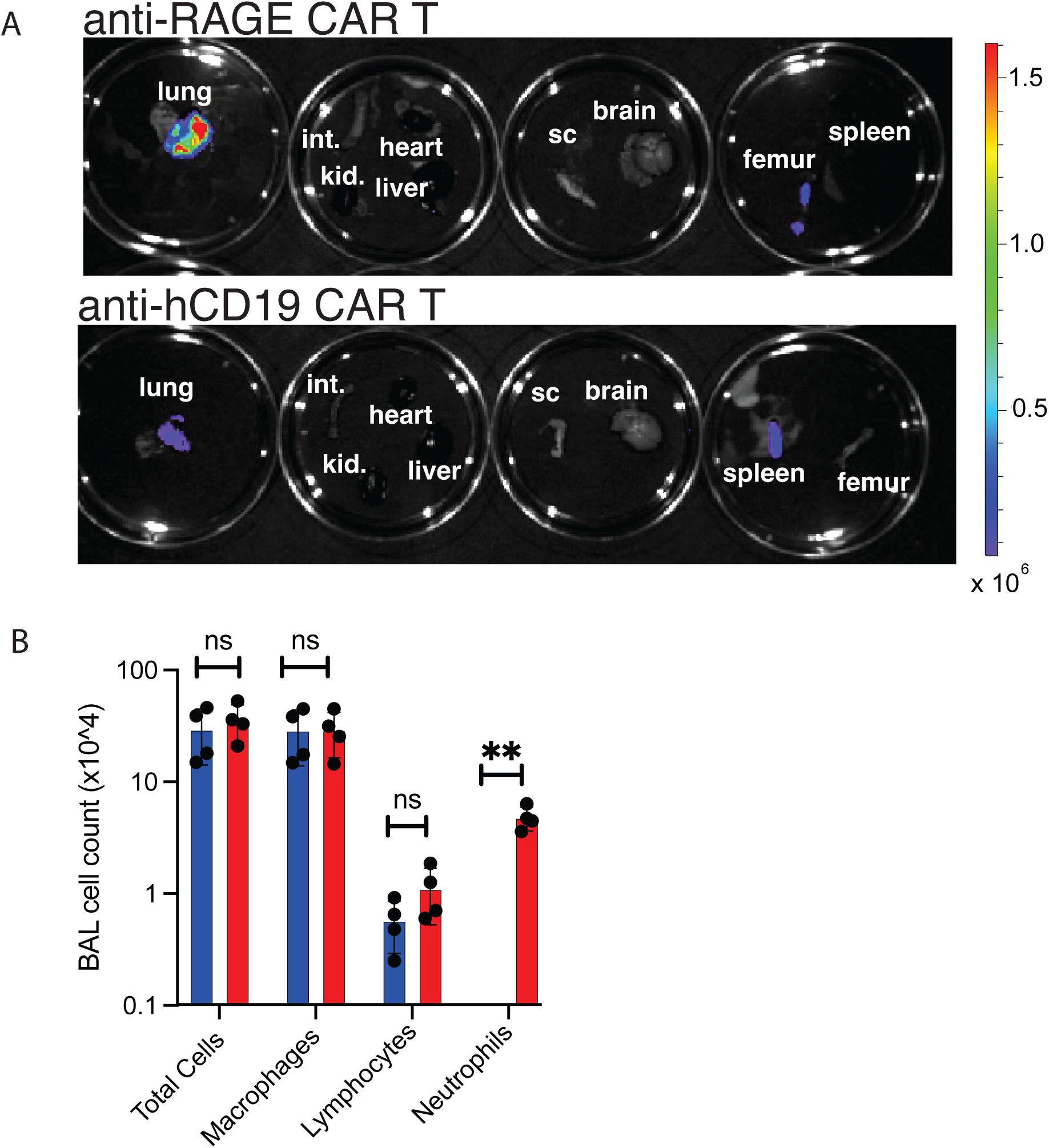
Anti-RAGE CAR traffics to the lung and causes alveolar leak. A) Bioluminescence of separate organs from mice 4 days after receiving either anti-RAGE CAR (top) or anti-human CD19 CAR (bottom) showing brightest signal in lung of anti-RAGE CAR mouse. B) Bronchoalveolar lavage fluid from lungs of mice from each group with cell counts for individual immune cell types. Healthy lungs have no neutrophils in lavage fluid. Increased neutrophils in anti-RAGE CAR group suggests inflammation leading to alveolar damage in this group.

**Supplementary Figure 3.**
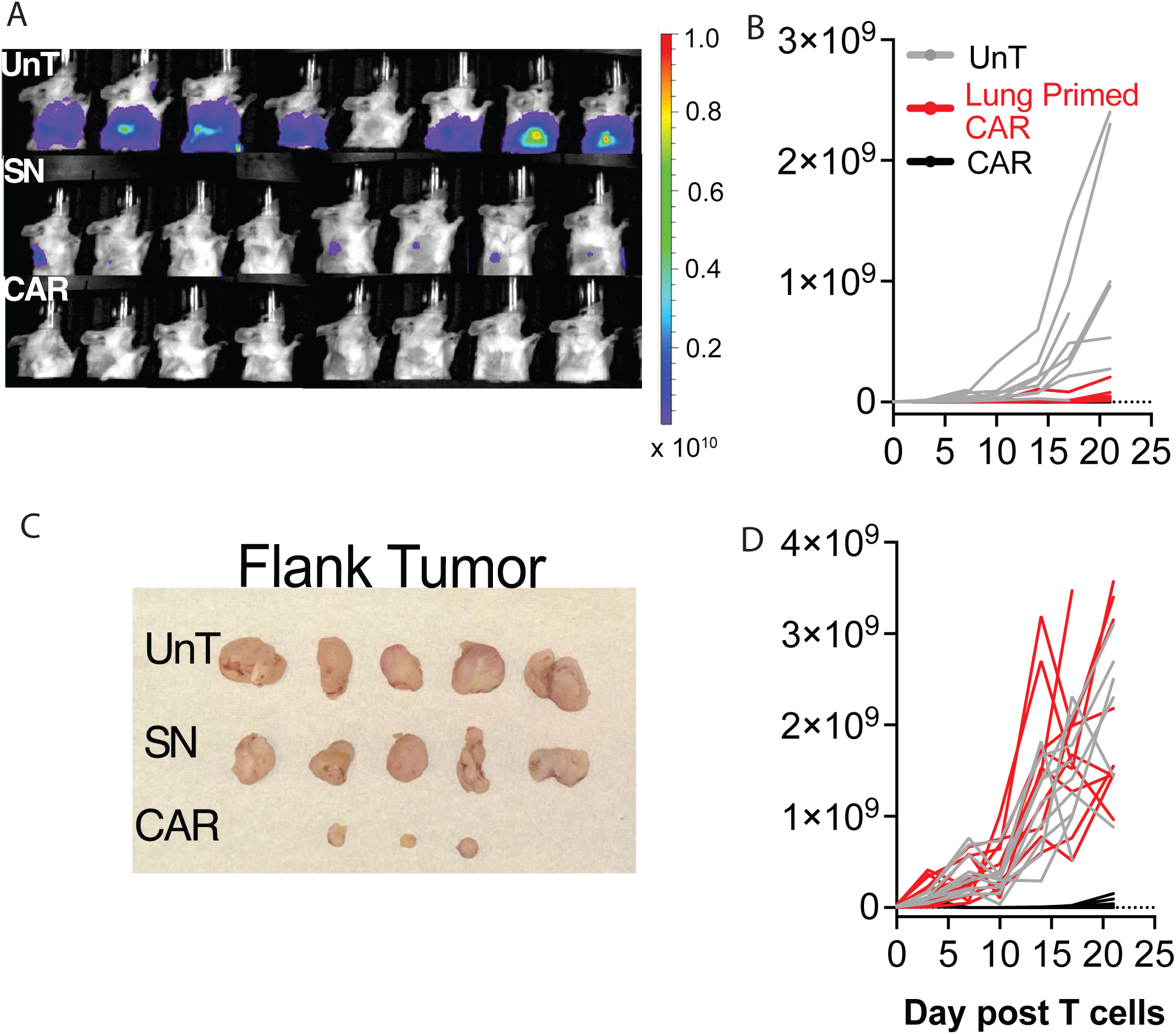
anti-RAGE SynNotch → CD19 CAR enables CD19 expressing tumor clearance specifically in the lung. A) Bioluminescent images processed to generate graphs (for lung fields, mice were placed right side up and bottom half of mice was covered such that the flank signal did not interfere with the lung signal. B) Data from individual mice represented in Figure 4F (lung). C) Images of flank tumors after experiment conclusion showing persistence of tumors in UnT and SN groups. D) Data from individual mice represented in Figure 4F (flank).

## Notes

### Competing Interest Statement

The authors have declared no competing interest.

### Funding

- National Heart Lung and Blood Institute: 5F32HL172536-02
- Advanced Research Projects Agency-Health: D24AC00084
- Marcus Program in Precision Medicine

